# Sero-prevalence and risk factors of leptospirosis in commercial cattle herds of Rupandehi district, Nepal

**DOI:** 10.1101/2020.07.29.226464

**Authors:** Tulsi Ram Gompo, Sumit Jyoti, Sudikchya Pandit, Ram Chandra Sapkota, Aditi Pandey

## Abstract

Nepal has an agrarian-based economy where the livestock sector contributes about 12 percent to the national GDP. Rupandehi district is one of the largest dairy pocket areas in the nation, but the disease, such as leptospirosis, posed a negative impact on their production and productivity. Despite the economic importance of cattle leptospirosis, the disease is concerned for the people’s occupational risk and food safety. Although leptospirosis is a priority zoonosis in Nepal, the effective implementation of the control program lacks both in animal and human health. A cross-sectional study was conducted to estimate the prevalence and identify associated risk factors of cattle leptospirosis from March 2019 to April 2020. Altogether, 206 cattle herds located in all the sixteen local levels of the district were visited. The owners were interviewed to record the cattle management system’s information and their knowledge of the zoonotic diseases. A total of 383 cattle serum samples were collected, and an indirect antibody ELISA was performed to estimate the infection status of leptospirosis in the cattle herds. All the data were analyzed by Open epi and R software for the descriptive and analytical study. Bivariate and multivariable logistic regression was applied to assess the potential herd level and animal level risk factors. Out of seven potential herd-level risk factors, the purchased cattle herds (OR: 7.2, 95% CI: 1.24-136.5, p=0.025) and cattle with herd size >10 (OR: 14.92, 95% CI: 2.61-283.38, p=0.025) were identified as significant risk factors for leptospirosis. At the animal level, the cattle taken for grazing in pastureland accessed by the community dog was a significant risk factor (OR: 4.16, 95% CI: 1.13-14.06, p=0.034). Surprisingly, none of the farmers had heard about leptospirosis before. The outcome of this research could be an epidemiological insight in control of priority zoonosis to protect the livestock economy and reduce their negative impact on public health.

## Background

Leptospirosis is a bacterial zoonosis, caused by the infection of leptospira species, which has a worldwide importance [1]. The natural carrier of this bacteria are wild and domestic animals, rodents, and dogs [2]. Among the various serovars, *Leptospira* hardjo is common in cattle [3–4], and they are the maintenance host for this serovar [4]. In cattle, leptospirosis causes reduced conception rates and reproductive problems like abortions, stillbirths, and decreased milk production [2,5].

People get infection after direct contact with infected animals or indirect contact with the contaminated environments by their urinals [6]. The occupational risk of exposures is mainly among the veterinarians, farmers, slaughterhouse workers, hunters, animal shelter workers, and agricultural workers[3]. World Health Organization (WHO) has considered leptospirosis a neglected and re-emerging zoonotic disease for humans and animals [7,8].

Leptospirosis is one of the six priority zoonoses listed for Nepal [9]; however, it’s adequate surveillance and reporting is still lacking in Nepal [10,11]. With the first report of human leptospirosis in 1981, numerous leptospirosis cases were reported among human and animal populations [12–15]. Most of these studies supported that the source of infection in humans is due to animal contact [13,14]. However, the actual socio-dynamic transmission of leptospirosis between the animals and the disease’s economic burden in both animals and humans was not well known.

Nepal’s one-third of the economy is based on agriculture, and the livestock sector singly contributes around 12% to the national GDP [15]. Cattle rearing is a traditional practice of Nepal [16], and the cattle ranks first among (7.2 million heads) all large ruminants in the country [17]. Moreover, the dairy industry has undergone rapid development since the last decades [18], but reproductive diseases such as leptospirosis in cattle have not been well assessed yet. The economic impact of the cattle production and its negative consequences on the livelihood of the poor and the marginal farmers is humongous [19].

There are only a few research articles and documentation about animal leptospirosis in cattle. According to research conducted in the Bhaktapur district of Nepal, the seroprevalence of *Leptospira interrogans* serovar hardjo was estimated at 5.11% (9/176) [20]. Another study over the various ecozones conducted in large ruminants of Nepal, the seroprevalence of *Leptospira* hardjo was 3.75% (12/320) during the pre-monsoon period and 6.88% (22/320) in post-monsoon period [21]. Similarly, a research conducted in infertile cattle and buffalo of western hills of Nepal showed the seropositivity of 11%(12/114) in cattle and 5.5% (5/86) in buffaloes [22].

This study aimed to identify the herd level and animal level risk factors that could contribute to the development of leptospirosis in cattle farming. The various risk factors associated with detecting leptospirosis in bovine leptospirosis were selected based on the literature review and expert’s opinion [23, 30]. The assessment of these risk factors at animal and herd levels could improve our understanding of leptospirosis, as the priority zoonosis could contribute to the development of control and contingency plan for it. It also attempts to assess farmers’ knowledge regarding the leptospirosis and make them aware of being protected during their livestock handling.

## Material and Methods

### Study sites

This study was conducted in the Rupandehi district located in the southwestern region of Nepal (Fig 1). The district has one of the highest number of dairy cattle with around 442 cattle farms [31], and about one hundred thousand of cattle population [17] within the total area of 530 square miles [15]. The district shares the border with neighboring nation India and has an international border quarantine where the importation of livestock exists between India and Nepal [32]. There is also a risk of disease spread due to the informal livestock trade and cattle’s internal movement between other adjacent districts.

**Fig 1:**
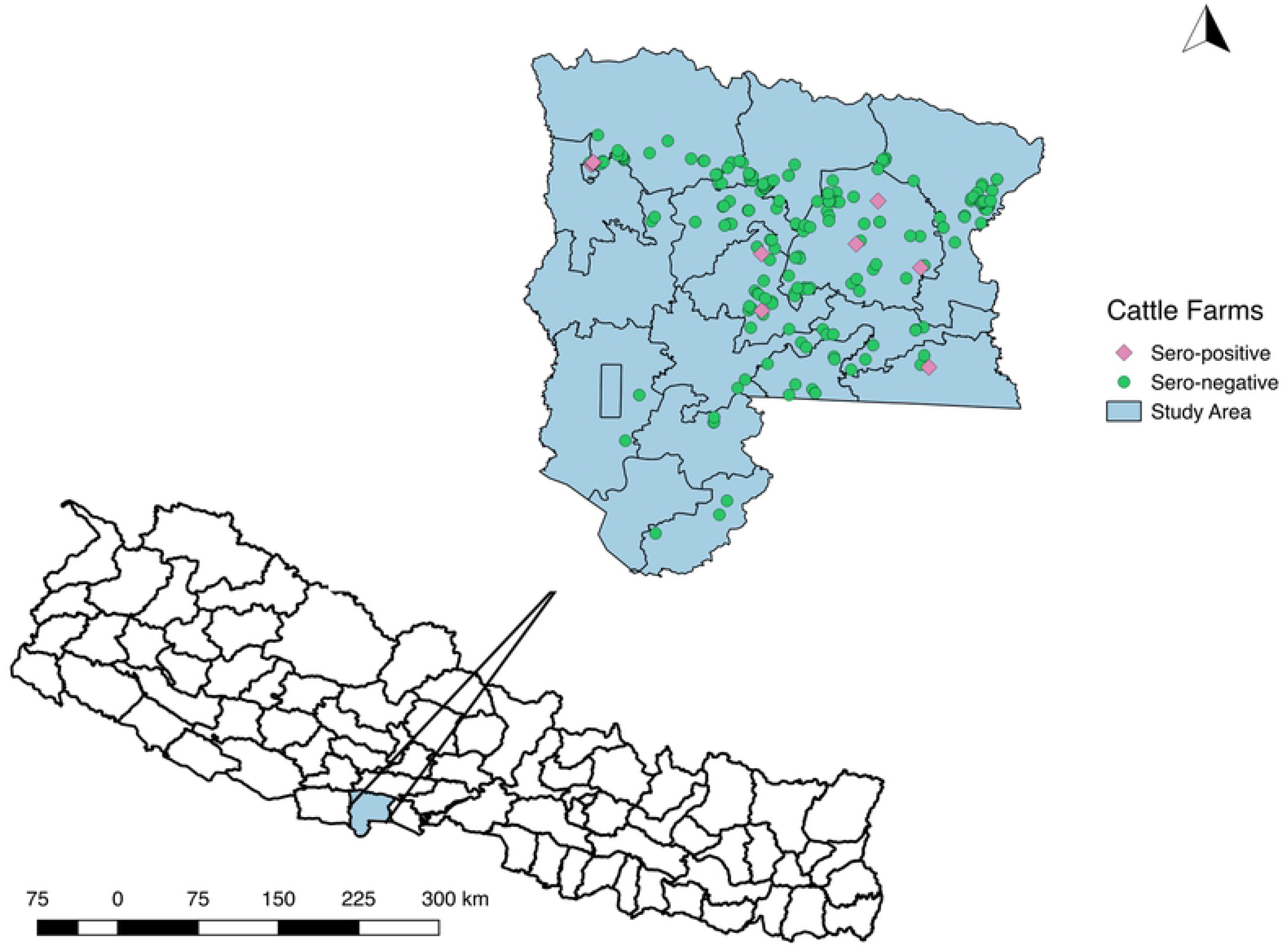
A map of Nepal, with the study district highlighted, including the seropositive and seronegative cattle farms.

### Study design

A cross-sectional study was conducted on commercial cattle herds of Rupandehi district between March 2019 to April 2020. The semi-structured questionnaire was used to collect the information on management status, biosecurity conditions, animal movement process of each herd, and owner’s knowledge of the principle zoonotic diseases such as leptospirosis, toxoplasmosis, and brucellosis. The questionnaire was initially designed in English and later translated to the local Nepali language (S1 file). A set of survey was pretested on ten farms as the pilot study. The response obtained from the owners were coded and analyzed. The response rate and clarity of the reaction were assessed to refine the survey questions. Next, blood samples were collected on the same day after interviewing the farm owners. The written consent from the owners was obtained during the questionnaire and sample collection processes.

### Sampling and sample size calculation

The sampling process consisted of two stages: first, herd-level sampling was done from total 442 commercial herds located in all sixteen local levels (municipalities) in the districts. Secondly, the propionate number of animals was selected from each herd giving a total individual sample size [33].

The sample size was determined using an open-source epidemiological software **“** OpenEpi” version 3.01 [34]. For selcting total herds, the expected prevalence was set as 50% with 5% desired precision at a 95% level of confidence that maximized the number of herd selection. The formula is based on:

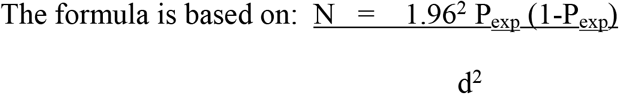

where N= sample size, Pexp = expected prevalence, and d = absolute precision [35].

The total number (N) of herd produced by this method is (N= 206). The same process as above, was repeated to calculate the total number of cattle to be sampled, keeping entire animal populations as 98,384 [17]. The total individual animal (n) to be sampled was (n= 383) samples. So, altogether 383 blood samples were collected from 206 selected herds located in the district.

### Sample transport and laboratory testing of samples

The tubes containing blood were allowed to remain overnight at room temperature to facilitate the serum separation, and the separated serum samples were transferred to a 2 ml Eppendorf tube. The collected serum samples were stored −20° Celsius and transported to Central Veterinary Laboratory for further laboratory analysis. The indirect ELISA against leptospirosis antibody was performed using the kit brought from Prionics Lelystad B.V., the Netherlands. The process was performed according to the protocols provided within the kit (citation). The reading of the ELISA plates was performed by the ELISA reader (“Multiskan™ FC Microplate Photometer”) at optical density (OD) 450nm within 15 minutes. The readings were interpreted with the software and protocol provided within the kits to determine the number of *L.* hardjo seropositive samples.

### Test result validation and interpretation criteria

After the measurement of optical density, the corrected OD value is calculated by subtracting the OD value of reference (controls) and test sera (samples) with the mean OD value of the blanks (ELISA buffer). Now, the percentage positivity (PP) of reference sera and the test sera were calculated as follows.

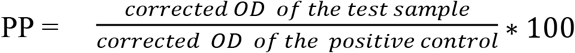

The result was validated if

i. The mean OD of the blanks was < 0.150
ii. The corrected OD of positive control was ≥1.000
iii. The mean PP of negative control was < 20
iv. The mean PP of weak positive control was between 20 to 60

For each sample, the cut off for percentage positivity for the test sera was determined as follows: PP < 20 was recorded negative, PP between 20-45 was classified as inconclusive, and PP ≥ 45 was considered positive for *L*. hardjo antibodies.

### Statistical Analysis

First, the raw data were entered manually on MS excel® spreadsheet from the paper-based questionnaire. After cleaning the data, the excel file was converted to a csv file and analyzed by R version 3.6.1 (R core Team 2019). The descriptive data analysis was performed to calculate the proportion of the seropositive herds with at least an animal positive to L. hardjo antibodies in it, and the percentage of seropositive animals among all herds. The true herd-level prevalence and the animal level prevalence were calculated by the use of open-source software Epitools by Ausvet [36]. The estimated sensitivity (Se) and the specificity (Sp) of the indirect antibody ELISA were estimated to be 96.4% and (Sp) 96.7%), respectively [37].

The continuous variables (herd size, age category, parity number) and an ordinal variable (body condition score) were converted to binary categorical variables using the quartiles (median as cut off) to avoid the problem of linearity [38]. The association between the categorical variables were assessed by Pearson’s Chi-square test. Highly related variables were assessed, and the biologically plausible variables were retained in the model for further analysis. For instance, there was a high correlation between the age category and the parity number and the cattle with common grazing ground and cattle grazing in community dog access. We believe that age could contribute to more critical information, and the community dog was a biologically the reservoir of leptospirosis, and these two variables were retained [39–43]. The reaming variables, the parity number and the common grazing ground, were ultimately dropped from the final model.

Bivariate analysis was performed to measure the association of each categorical variable with the outcome “seropositivity to leptospirosis” (Table 1 and 2). The variables with the significance level p≤0.2 following the bivariate analysis and those with the biological plausibility were manually entered in the final multivariable model [44]. A backward stepwise variable selection was used to add the variable with the lowest p-value to construct a final model with a significance level of p ≤ .05. Any variables with p-value <0.05 were considered statistically significant and considered the risk factors. (Table 3 and Table 4).

**Table 1:** Herd level factors potentially associated with seropositive to leptospira by univariate logistic regression analysis

| Determinants | Factor level | Total number of herds (n=206) | Positive farms | Negative farms | OR | 95% CI | P- value |
| --- | --- | --- | --- | --- | --- | --- | --- |
| **Herd Characteristics** |  |  |  |  |  |  |  |
| Cattle origin | Purchased | 125 | 9 | 116 | 6.21 | (1.13-115.6) | 0.09 |
|  | Home breed | 81 | 1 | 80 |  |  |  |
| Herd Size<br>(median=10) | >10 | 93 | 9 | 84 | 12.00 | (2.19-223.44) | 0.02* |
|  | <=10 | 113 | 1 | 112 |  |  |  |
| **Management system** |  |  |  |  |  |  |  |
| Herd structure | Head to head | 71 | 6 | 65 | 3.29 | (0.84-16.02) | 0.10<sup>a</sup> |
|  | Tail to tail | 25 | 1 | 24 | 1.49 | (0.07-12.19) | 0.74 |
|  | Single | 110 | 3 | 107 | Ref |  |  |
| Bedding type | Mat | 22 | 1 | 21 | 1.81 | (0.08,16.1) | 0.62 |
|  | None | 144 | 3 | 141 | 0.58 | (0.1,3.2) | 0.51 |
|  | Saw dust | 96 | 4 | 92 | 0.92 | (0.16-5.2) | 0.93 |
|  | Straw | 105 | 6 | 99 | Ref |  |  |
| Source of water | Tap | 116 | 6 | 110 | 1.17 | (0.32-4.71) | 0.81 |
|  | Well | 90 | 4 | 86 | Ref |  |  |
| Feeding system | Occasional grazing | 13 | 1 | 12 | 1.704 | (0.09-10.25) | 0.63 |
|  | Stall feeding | 193 | 9 | 184 | Ref |  |  |
| Breeding type | AI | 175 | 9 | 166 | 1.70 | (0.09-10.25) | 0.65 |
|  | Natural<br>Breeding | 31 | 1 | 30 | Ref |  |  |
| Cat feces seen around<br>farms | Yes | 72 | 5 | 67 | 1.94 | (0.52-7.20) | 0.31 |
|  | No | 134 | 5 | 129 | Ref |  |  |
| Feed contamination<br>with the soil | Yes | 127 | 8 | 119 | 2.59 | (0.63-17.45) | 0.20<sup>a</sup> |
|  | No | 79 | 2 | 77 | Ref |  |  |
| Rodents seen around<br>farms | Yes | 139 | 9 | 130 | 4.57 | (0.83-85.16) | 0.09 |
|  | No | 67 | 1 | 66 | Ref |  |  |
| Owners worked on rice<br>field with bare feet | Yes | 148 | 9 | 139 | 3.691 | (0.67,68.84) | 0.15<sup>a</sup> |
|  | No | 58 | 1 | 57 | Ref |  |  |

a: these non-significant variables were entered in the final model with cut off criteria of $p \leq 0.2$ , including the significant variables.
\*P value <0.05 are the significant variables.

**Table 2:**
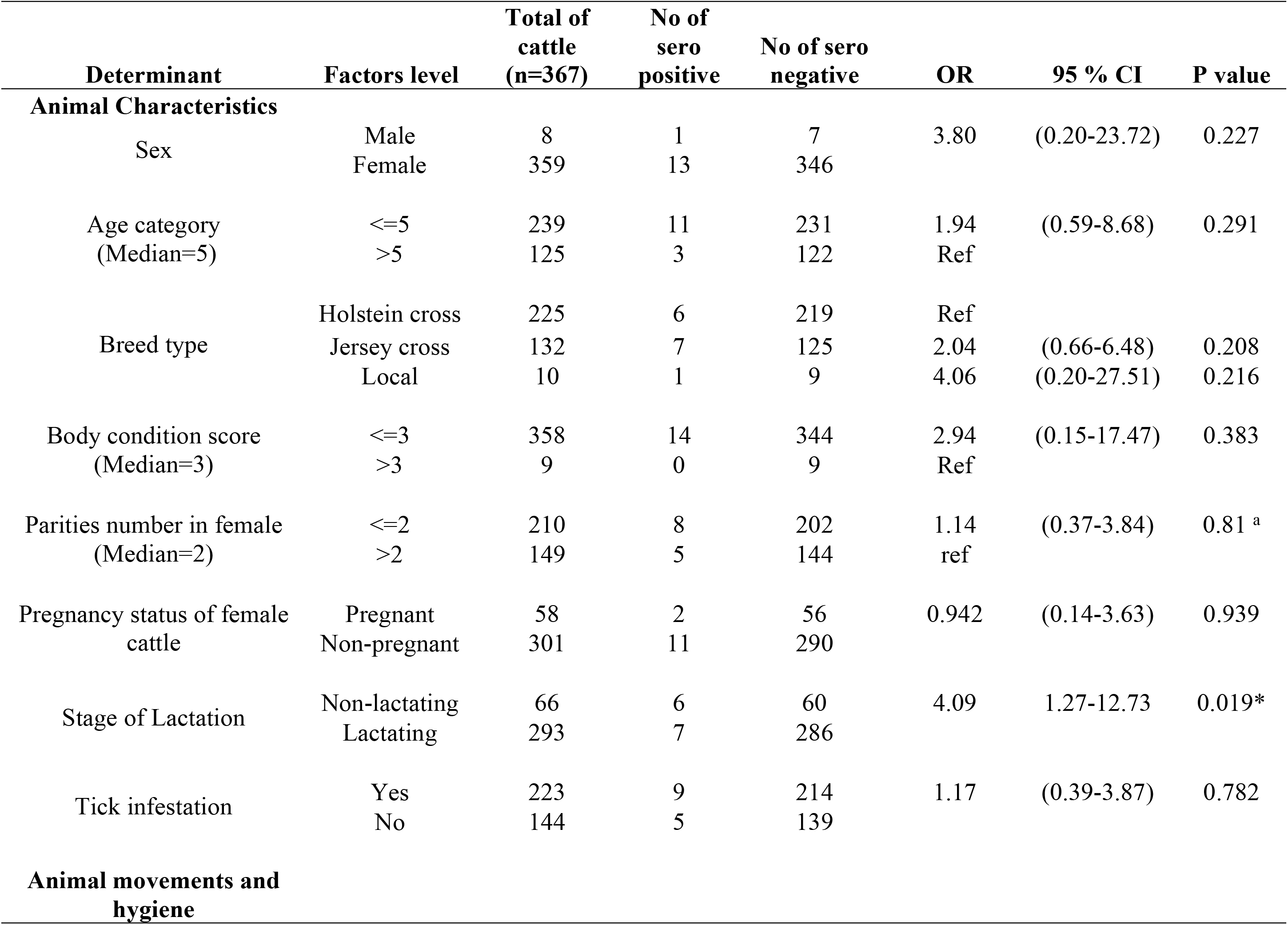
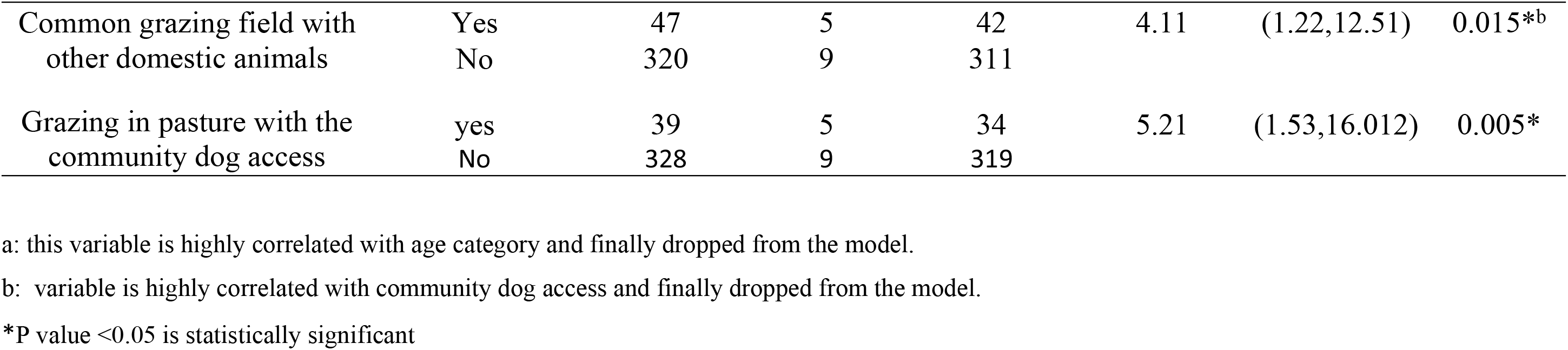
Animal level factors potentially associated with seropositive to leptospira by univariate logistic regression analysis

**Table 1:** Herd level and animals risk factors associated with leptospirosis infection on commercial dairy farms by multivariable logistic regression analysis.

| Determinants | Response level | Adjusted OR | 95%CI | p value |
| --- | --- | --- | --- | --- |
| **i. Herd level risk factors (n=206)** |  |  |  |  |
| Cattle origin | Purchased<br>Homebred | 7.2 | (1.24,136.5) | 0.025* |
| Herd size | >10<br><=10 | 14.92 | (2.61, 283.38) | 0.001* |
| Owners working bare foot on rice fields | Yes<br>No | 5.67 | (0.96, 108.75) | 0.057 |
| Rodents seen around kitchen and farms | Yes<br>No | 3.93 | (1.53-75.72) | 0.149 |
| **ii. Animal Level risk factors (n = 367)** |  |  |  |  |
| Grazing in pasture with the community dog access | Yes<br>No | 4.16 | (1.13- 14.06) | 0.034* |
| Production status of cattle | Lactating<br>Non-lactating | 4.09 | (0.84-9.59) | 0.089 |

CI: confidential intervals;
OR: odds ratios;
\*P-value <0.05

The multicollinearity among the independent variables was checked for variance inflation factors (VIF) by “vif” function in R, and variables with “vif” coefficient >10 were considered high. None of the independent variables in the analyses showed multicollinearity.

The potential confounding effects of the variables were checked with the changes in the point estimates of the variables that remain in the model [43–45]. Any changes in the coefficient with >20% were included in the final model.

Alternatively, a full model was built in the first place, and the appropriate model was selected based on the automatic stepwise Akaike’s Information Criteria (AIC) selection process (both backward and forward direction). The initial manually derived models were compared to the automated derived models, but all of the approaches ended up the same set of variables on the final model (Table 3 and Table 4). The fitness of the final models was assessed by Hosmer-Lemeshow goodness of fit function (hoslem.test) in R, and they were fit.

### Ethical statement

Ethical statements for animal subjects were approved by Nepal Veterinary Council (NVC), Ref. No 179 (TG)/2076.77, the official statutory body of animal health. A written consent of form from the cattle owner were obtained during the interview. The experienced and registered veterinarian handled all the animals during animals’ sampling, taking account of minimal pain during the blood collection. The owners also requested to provide the test results of their cattle herds through confidential means.

## Results

### Descriptive study

Of the total 206 herds selected, the median herd size was 10 with a range of one to 200 cattle. The median age of the cattle in the herd was five years old. Of the total 206 herd, the composition of home bred and purchased animals was 60.68% (125/206) and 39.32% (81/206). The majority, 93.69% (193/206), of the herds, were based on stall-feeding system, and only 6.31% (13/206) of them were taken for occasional grazing. About eighty-five percentage (175/206) of the farms applied an artificial insemination process for breeding their animals, and the remaining 15.04% (31/206) of the herds used to apply the natural mating system. Interestingly, 22% (67/300) of the farmers used to work both in the rice field and the cattle rearing. However, none of the farmers had heard about the leptospirosis before, and none had any information to prevent transmission of that diseases from their livestock.

### Herd level seroprevalence and potential risk factors

Of the total 206 herds tested against leptospirosis, 10 (4.85%,95%CI: 1.92-7.79) herds had at least one seropositive animal. The potential herd level variables were classified broadly into two categories: i) Herd characteristics and Herd management system [24–25, 29, 39, 43, 46].

### Animal level seroprevalence and potential risk factors

Of the 383 blood samples collected, 16 samples produced hemolysis, and they were discarded from testing. Of the total 367 serum samples, 14 (3.81 %, 95%CI: 2.29, 6.3) were tested positive to *L.* hardjo. The potential animal level risk factors selected were: i) animal characteristics such as sex, age, breed, body condition score, parity number, pregnancy status, stage of lactation, tick infestation ii) animal movement patterns such as taking animals in the grazing field and pasture access by the community dogs [24,29,43,47].

### Univariate regression analysis for herd and animal level risk factors

There were 11 herd-level risk factors included in the univariate analysis, as depicted in Table 1. The purchased cattle herds had higher odds of detecting Leptospira hardjo (OR: 6.21, 95% CI: (1.13-115.6, p=0.09) compared to the home-bred herds, but the finding was only borderline significant. The cattle herds with median herd size of >10 were 12 times more likely to be seropositive compared to those with a herd size of ≤ 10 (OR: 12.00, 95% CI: 2.19-223.44, p=0.02). Among the potential herd-level risk factors, the variables with the cut-off values of p≤ 0.20 were qualified to enter in the final multivariable model. Such variables were cattle herd origin, herd size, herd structure, herd fed with soil contaminated feed, presence of rodents on farms, and the herd whose owners worked in the rice field with bare feet (Table 1).

Ten potential animal level risk factors were selected and classified under broad category: animal characteristics and animal movement pattern, as shown in Table 2. In the univariate analysis, the non-lactating cattle were four times (OR: 4.09, 95% CI: 1.27-12.73, p=0.019), more likely to be seropositive with leptospirosis compared to lactating cattle. The cattle taken for common grazing in the fields were four times more likely to be seropositive to leptospira (OR: 4.11, 95% CI: 1.22,12.51, p=0.015) compared to animals that were stall-fed on farms. The animals grazed on the field with the community dog access had significantly higher odds of leptospira seropositivity (OR: 5.21, 95% CI: 1.53,16.012, p=0.005). However, none of the other animal level risk factors were associated with the seropositive to leptospirosis among cattle.

The variables “parity number” and “common grazing ground” were dropped because they were highly associated with “age category” and “pasture grazing with community dog access,” respectively. The set of variables such as the breed of cattle, animals’ parities, history of repeat breeding, stage of lactations, and grazing in pasture with the community dog access were selected for multivariable analysis.

### Multivariable logistic regression Analysis for herd and animal level risk factors

Multivariable logistic regression analyses were separately conducted for a group of the herd-level and the animal-level risk factors. Seven variables from the set of herd-level and four variables from the animal level risk factors were qualified to enter the final multivariable models based on cut off criteria of p-value ≤ 0.2.

In the final herd level multivariable model, the two herd-level risk factors: cattle origin and herd size were associated with seropositivity to leptospirosis (Table 3). The purchased cattle herds had significantly higher odds of detecting leptospirosis (OR: 7.2, 95% CI: 1.24-136.5, p=0.025) compared to home-bred cattle herds. Cattle herds with herd size >10 animals were about fifteen times more likely to seropositive for leptospirosis (OR: 14.92, 95% CI: 2.61-283.38, p=0.025) than herd size of ≤ 10 animals.

In the final animal level multivariable model (Table 3), the animals taken for grazing in the pastureland with access to community dogs had significantly higher odds of seropositive to leptospirosis (OR: 4.16, 95% CI: 1.13-14.06, p=0.034). The cattle herd owners who worked in the rice fields showed an increased likelihood of leptospira seropositivity in their cattle herds (borderline significance; OR: 5.67, 95% CI: 0.96-108.75, p=0.057). Non-lactating animals that was a risk factor in the univariable analysis was not a risk for leptospira seropositivity in the multivariable model (OR: 4.09, 95% CI: 0.84-9.59, p=0.089).

## Discussion

We estimated that the herd level and animal level seroprevalence of leptospirosis as 4.85% (10/206) and 3.81 % (14/367), respectively, among the cattle herds in Rupandehi district of Nepal. There were some previous regional studies in Nepal, that estimated 3.75 % (12/320) to 11% (12/114) of the leptospirosis in cattle population[21,22]. Nevertheless, there might be either overestimation or underestimation of the leptospirosis prevalence due to differences in epidemiological design, sample size, and geographical diversity. Probably, this is the first risk factor study of leptospirosis among cattle in Nepal to our knowledge.

The purchased herds were more likely to detect leptospirosis than the homebred herds in our study. It is epidemiologically justifiable that the herds that moved during selling are more likely to contract the disease [12, 48–49].

Also, the Rupandehi district of Nepal is located in the trade transit with India and neighboring provinces, so the cattle herd, when moved from one destination to another, have higher exposures compared to the confined herds.

Our findings show that the cattle herds of >10 animals in the herd was a risk factor. The bigger herd size is the risk factor for leptospirosis, and many of the studies have already proved that [25,29]. The high stocking density in the bigger herd size farms had a higher risk of getting the diseases due to frequent selling or purchasing of animals, and the contact between infectious and susceptible animals takes place more often [50].

The cattle herds owned by the people who worked in rice fields with bare feet were at borderline risk than herds whose owners had another occupation (p<0.05). It was because most of the cattle owners in Nepal were the farmers, and they were more likely to work in the rice field to get straw for the cattle. Moreover, the possible environmental contamination might be a source of infection in the herds. The above finding was similar to the study by Alavi et al. [51] in southwestern Iran, and Sejvar et al. [52] in Malaysia. The owners who used to work in the rice fields posed risks to their cattle and themselves unless they apply the personal protective measures.

Though rodents are the carriers for leptospirosis [2], the association was not statistically significant in the multivariable analysis. However, rodents should be controlled in the farms because they could contribute to transmission of leptospira organisms.

Our findings showed that the cattle taken in the pasture with community dog access had significantly higher odds of leptospirosis. In Nepal, there are a significant number of community dogs in the cities and village areas. These dogs frequently move in search of food and defecate openly in the pasture that leads to the contamination of grazing ground. The above statement is supported by the findings in Brazil by Fávero et al. [43], who also demonstrated that the dog in the pasture was associated with detecting leptospira antibodies in cattle. This study findings might advocate the local governments to formulate plans in managing the population of community dogs around the livestock farming area.

However, our study also had some limitations. Some cattle farm owners do not remain on the farms, and the information was obtained from the less educated farmworkers, and their level of understanding may not represent the idea of the all cattle owners in general. There could be potential recall bias in obtaining the animals’ information as most of them had not maintained the excellent record-keeping on the farms.

## Conclusions

This research estimated the burden and identified the risk factors of cattle leptospirosis in the commercial cattle herds that provided an epidemiological insight into animal leptospirosis in Nepal. Though leptospirosis has been documented as priority zoonoses by the government, the effective implementation of the surveillance program still lacks in the nation. The outcome of this research might be useful for effective program planning in the prevention and control of leptospirosis both in animal and human health. Ultimately, the livestock production and productivity should be enhanced, food safety for the public must be assured, and farmworkers’ occupational safety should be ensured. We suggest further nationwide research and application of control measures through one health approach to reduce the incidence of zoonotic diseases like leptospirosis in Nepal.

## Supporting information

S1: Questionnaire used for the study

## Acknowledgement

We acknowledge Dr. Diker Dev Bhatta, Chief of Central Veterinary Laboratory for arranging the resources for testing, and Dr. Luna Gongal for assisting in lab works. We want to thank Dr. Sangeeta Rao of Colorado State University providing suggestions, and revision on the manuscript. Lastly, we want to acknowledge all the helping hands and the cattle farmers of Rupandehi district who cooperated to participate in our study.

## Authors contribution

TRG: Conceived the study design, conducted the research, performed lab, analyzed data and wrote the paper, revised draft.SJ: assisted in data and sample collection, lab testing and literature review. SP and AP: Assisted in data and sample collection. RC: assisted in lab testing.

## Conflict of interest

Authors declare no conflict of interest with this research.

## Funding

Authors receive no special funding for this work.

## References

1. Mohammed H, Nozha C, Hakim K, Abdelaziz F. LEPTOSPIRA: Morphology, Classification and Pathogenesis. J Bacteriol Parasitol. 2011;02. doi:10.4172/2155-9597.1000120

2. Adler B, de la Peña Moctezuma A. Leptospira and leptospirosis. Vet Microbiol. 2010;140: 287–296. doi:10.1016/j.vetmic.2009.03.012

3. Levett PN. Leptospirosis. Clin Microbiol Rev. 2001;14: 296–326. doi:10.1128/CMR.14.2.296-326.2001

4. Grooms DL. Reproductive losses caused by bovine viral diarrhea virus and leptospirosis. Theriogenology. 2006;66: 624–628. doi:10.1016/j.theriogenology.2006.04.016

5. Dhaliwal GS, Murray RD, Dobson H, Montgomery J, Ellis WA. Reduced conception rates in dairy cattle associated with serological evidence of Leptospira interrogans serovar hardjo infection. Vet Rec. 1996;139: 110–114. doi:10.1136/vr.139.5.110

6. Haake DA, Levett PN. Leptospirosis in Humans. In: Adler B, editor. Leptospira and Leptospirosis. Berlin, Heidelberg: Springer Berlin Heidelberg; 2015. pp. 65–97. doi:10.1007/978-3-662-45059-8_5

7. Vijayachari P, Sugunan AP, Shriram AN. Leptospirosis: an emerging global public health problem. J Biosci. 2008;33: 557–569. doi:10.1007/s12038-008-0074-z

8. World Health Organisation. Leptospirosis Burden Epidemiology Reference Group (LERG). 2010. Available: https://www.who.int/zoonoses/diseases/lerg/en/

9. MInistry of Health and Population. Zoonois Control Program. 2012. Available: https://www.mohp.gov.np/eng/program/communicable-disease/zoonotic-disease-control-programme

10. Bhattachan B, Bhattachan A, Sherchan JB, Dhoubhadel BG, Sherchand JB. Leptospirosis: An Emerging Infectious Disease in Nepal. 2016. Available: https://www.researchgate.net/publication/315496847_Leptospirosis_An_Emerging_Infectious_Disease_in_Nepal

11. Dahal KP, Sharma S, Sherchand JB, Upadhyay BP, Bhatta DR. Detection of Anti-*Leptospira* IgM Antibody in Serum Samples of Suspected Patients Visiting National Public Health Laboratory, Teku, Kathmandu. Int J Microbiol. 2016;2016: 1–4. doi:10.1155/2016/7286918

12. Myint KSA, Murray CK, Scott RM, Shrestha MP, Mammen MP, Shrestha SK, et al. Incidence of leptospirosis in a select population in Nepal. Trans R Soc Trop Med Hyg. 2010;104: 551–555. doi:10.1016/j.trstmh.2010.04.001

13. Dahal KP, Sharma S, Sherchand JB, Upadhyay BP, Bhatta DR. Detection of Anti-*Leptospira* IgM Antibody in Serum Samples of Suspected Patients Visiting National Public Health Laboratory, Teku, Kathmandu. Int J Microbiol. 2016;2016: 1–4. doi:10.1155/2016/7286918

14. Shrestha R, McKenzie JS, Gautam M, Adhikary R, Pandey K, Koirala P, et al. Determinants of clinical leptospirosis in Nepal. Zoonoses Public Health. 2018;65: 972–983. doi:10.1111/zph.12516

15. Central Bureau of Statistics, Nepal. Nepal Census Info 2011. Nepal; 2011. Available: http://dataforall.org/dashboard/nepalcensus/

16. Cow farm in Nepal. 2020. Available: http://www.agricultureinnepal.com/cow-farm

17. Ministry of Livestock Development. LIVESTOCK STATISTICS OF NEPAL. Kathmandu, Nepal; 2017. Available: https://nepalindata.com/resource/livestock-statistics-of-nepal-2017--/

18. Food and Agriculture Organization of the United Nations. Dairy Sector Study of Nepal. Nepal; 2010. Available: http://nepalagritech.com.np/wp-content/uploads/2016/10/Dairy-Sector-in-Nepal-FAO.pdf

19. FAO,. ocio-economic consequences for poor livestock farmers of animal diseases and VPH problems. Rome; 2002. Available: http://www.fao.org/3/Y3542E/y3542e00.htm#Contents

20. Rawal G, Shrestha D. Sero-Detection of Leptospira hardjo in Cattle of Bhaktapur District of Nepal. Int J Appl Sci Biotechnol. 2019;7: 378–381. doi:10.3126/ijasbt.v7i3.25713

21. Khanal et al. Detection of Antibodies Against Leptospira hardjo in Large Ruminants of Nepal. ACTA Sci Agric. 2018;2.

22. Joshi, H. D., Joshi, B. R. Detection of Leptospira hardjo antibodies in infertile cattle and buffaloes in the western hills of Nepal. Nepal Agric Res Counc Anim Health Res Div. 2000;15. Available: https://www.cabdirect.org/cabdirect/abstract/20023063242

23. Lilenbaum W, Souza GN. Factors associated with bovine leptospirosis in Rio de Janeiro, Brazil. Res Vet Sci. 2003;75: 249–251. doi:10.1016/S0034-5288(03)00114-0

24. Schoonman L, Swai ES. Herd- and animal-level risk factors for bovine leptospirosis in Tanga region of Tanzania. Trop Anim Health Prod. 2010;42: 1565–1572. doi:10.1007/s11250-010-9607-1

25. Ryan EG, Leonard N, O’Grady L, Doherty ML, More SJ. Herd-level risk factors associated with Leptospira Hardjo seroprevalence in Beef/Suckler herds in the Republic of Ireland. Ir Vet J. 2012;65: 6. doi:10.1186/2046-0481-65-6

26. Dreyfus, Anou. Leptospirosis in humans and pastoral livestock in New Zealand : a thesis presented in partial fulfilment of the requirements for the doctoral degree of Doctor of Philosophy at Massey University. 2013. Available: http://hdl.handle.net/10179/4870

27. Loan HK, Van Cuong N, Takhampunya R, Kiet BT, Campbell J, Them LN, et al. How Important Are Rats As Vectors of Leptospirosis in the Mekong Delta of Vietnam? Vector-Borne Zoonotic Dis. 2015;15: 56–64. doi:10.1089/vbz.2014.1613

28. Schuller S, Francey T, Hartmann K, Hugonnard M, Kohn B, Nally JE, et al. European consensus statement on leptospirosis in dogs and cats. J Small Anim Pract. 2015;56: 159–179. doi:10.1111/jsap.12328

29. Miyama T, Watanabe E, Ogata Y, Urushiyama Y, Kawahara N, Makita K. Herd-level risk factors associated with Leptospira Hardjo infection in dairy herds in the southern Tohoku, Japan. Prev Vet Med. 2018;149: 15–20. doi:10.1016/j.prevetmed.2017.11.008

30. Olmo L, Reichel MP, Nampanya S, Khounsy S, Wahl LC, Clark BA, et al. Risk factors for Neospora caninum, bovine viral diarrhoea virus, and Leptospira interrogans serovar Hardjo infection in smallholder cattle and buffalo in Lao PDR. Stevenson B, editor. PLOS ONE. 2019;14: e0220335. doi:10.1371/journal.pone.0220335

31. Veterinary Hospital and Livestock Service Center, Rupandehi. Annual Report. Nepal; 2019.

32. Central Animal Quarantie Office, Kathmandu. Animal Quarantine Monthly Report 2018. Nepal; 2018. Available: http://www.caqo.gov.np/uploads/files/5453250888.pdf

33. Skinner CJ. Probability Proportional to Size (PPS) Sampling. In: Balakrishnan N, Colton T, Everitt B, Piegorsch W, Ruggeri F, Teugels JL, editors. Wiley StatsRef: Statistics Reference Online. Chichester, UK: John Wiley & Sons, Ltd; 2016. pp. 1–5. doi:10.1002/9781118445112.stat03346.pub2

34. Dean AG, Sullivan KM, Soe MM. OpenEpi:Open Source Epidemiologic Statistics for Public Health. USA; 2013. Available: www.OpenEpi.com

35. Michael Thrusfield. Veterinary epidemiology. Fourth. Willey and Blackwell; 2018.

36. Sergeant, ESG. Epitools Epidemiological Calculators. Australia; 2018. Available: http://epitools.ausvet.com.au.

37. Lewis FI, Gunn GJ, Mckendrick IJ, Murray FM. Bayesian inference for within-herd prevalence of Leptospira interrogans serovar Hardjo using bulk milk antibody testing. Biostatistics. 2009;10: 719–728. doi:10.1093/biostatistics/kxp026

38. Turner EL, Dobson JE, Pocock SJ. Categorisation of continuous risk factors in epidemiological publications: a survey of current practice. Epidemiol Perspect Innov. 2010;7: 9. doi:10.1186/1742-5573-7-9

39. Boqvist S, Chau BL, Gunnarsson A, Olsson Engvall E, Vågsholm I, Magnusson U. Animal-and herd-level risk factors for leptospiral seropositivity among sows in the Mekong delta, Vietnam. Prev Vet Med. 2002;53: 233–245. doi:10.1016/S0167-5877(01)00263-X

40. Claus Anita, Van de Maele, Pasmans F, Gommeren K, Daminet S. Leptospirosis in dogs: a retrospective study of seven clinical cases in Belgium. VLAAMS DIERGENEESKUNDIG TIJDSCHRIFT. 77: 259–264. Available: http://hdl.handle.net/1854/LU-675967

41. van de Maele I, Claus A, Haesebrouck F, Daminet S. Leptospirosis in dogs: a review with emphasis on clinical aspects. Vet Rec. 2008;163: 409–413. doi:10.1136/vr.163.14.409

42. Ellis WA. Control of canine leptospirosis in Europe: time for a change? Vet Rec. 2010;167: 602–605. doi:10.1136/vr.c4965

43. Fávero JF, de Araújo HL, Lilenbaum W, Machado G, Tonin AA, Baldissera MD, et al. Bovine leptospirosis: Prevalence, associated risk factors for infection and their cause-effect relation. Microb Pathog. 2017;107: 149–154. doi:10.1016/j.micpath.2017.03.032

44. Dahoo I, Martin W, Stryhn H. Veterinary epidemiologic research. 2. ed., 3. print. Charlottetown, CA: VER Inc; 2014.

45. Hosmer DW, Lemeshow S. Applied logistic regression. New York; Toronto: John Wiley & Sons; 2005. Available: http://onlinelibrary.wiley.com/book/10.1002/0471722146

46. Wójcik-Fatla A, Zając V, Cisak E, Sroka J, Sawczyn A, Dutkiewicz J. Leptospirosis as a tick-borne disease? Detection of Leptospira spp. in Ixodes ricinus ticks in eastern Poland. Ann Agric Environ Med AAEM. 2012;19: 656–659.

47. Boqvist S, Chau BL, Gunnarsson A, Olsson Engvall E, Vågsholm I, Magnusson U. Animal- and herd-level risk factors for leptospiral seropositivity among sows in the Mekong delta, Vietnam. Prev Vet Med. 2002;53: 233–245. doi:10.1016/S0167-5877(01)00263-X

48. Oliveira FCS, Azevedo SS, Pinheiro SR, Batista CSA, Moraes ZM, Souza GO, et al. Fatores de risco para a leptospirose em fêmeas bovinas em idade reprodutiva no Estado da Bahia, Nordeste do Brasil. Pesqui Veterinária Bras. 2010;30: 398–402. doi:10.1590/S0100-736X2010000500004

49. Hashimoto VY, Dias JA, Spohr KAH, Silva MCP, Andrade MGB, Müller EE, et al. Prevalence and risk factors for Leptospira spp. in cattle herds in the south central region of Paraná state. Pesqui Veterinária Bras. 2012;32: 99–105. doi:10.1590/S0100-736X2012000200001

50. Yupiana Y, Vallée E, Wilson P, Weston JF, Benschop J, Collins-Emerson J, et al. On-farm risk factors associated with *Leptospira* shedding in New Zealand dairy cattle. Epidemiol Infect. 2020; 1–24. doi:10.1017/S095026882000103X

51. Alavi L, Alavi SM, Khoshkho MM. Risk Factors of Leptospirosis in Khuzestan, South West of Iran, 2012. Int J Enteric Pathog. 2013;1: 68–71. doi:10.17795/ijep15248

52. Sejvar J, Bancroft E, Winthrop K, Bettinger J, Bajani M, Bragg S, et al. Leptospirosis in “Eco-Challenge” Athletes, Malaysian Borneo, 2000. Emerg Infect Dis. 2003;9: 702–707. doi:10.3201/eid0906.020751

